# EZH2 inhibition enhances TRAIL responses in multiple myeloma but not in quiescent cells

**DOI:** 10.1101/2021.07.23.453217

**Authors:** Amal Arhoma, Jenny Southan, Andrew D Chantry, Sarah L Haywood-Small, Neil A Cross

**Affiliations:** Biomolecular Sciences Research Centre, Sheffield Hallam University, Sheffield, UK; Sheffield Teaching Hospitals NHS Foundation Trust; Mellanby Centre for Bone Research, Department of Oncology and Metabolism, University of Sheffield, UK

**Keywords:** Multiple Myeloma, TRAIL, Apoptosis, Histone Methyltransferase, EZH2, Quiescence

## Abstract

Multiple Myeloma is a plasma cell malignancy for which there is currently no cure, despite many novel therapies. TRAIL (Tumour necrosis factor-related apoptosis inducing ligand) is a promising anti-tumour agent although TRAIL-insensitive cells readily emerge when used as a single agent and its effects are limited due to many TRAIL-resistant cells which emerge soon after treatment. EZH2 is a H3K27 histone methyltransferase found to be overexpressed in many cancers including Multiple Myeloma. We tested the hypothesis that epigenetic reprogramming using the EZH2 inhibitor GSK343 would enhance TRAIL sensitivity, overcome TRAIL resistance, and target TRAIL-resistant quiescent cell populations. We show that GSK343 is a potent TRAIL sensitiser in TRAIL-sensitive RPMI 8226, NCI-H 929 and U266, and in TRAIL-resistant OPM-2 and JJN3, the latter showing very potent synergistic induction of apoptosis, primarily via caspase-8 activation but also via caspase-9. GSK343-enhancement of TRAIL responses was further enhanced in a 3D cell culture model of Multiple Myeloma in NCI-H 929 and U266. We show that in TRAIL-resistant sub-populations of Multiple Myeloma cells, GSK343 responses were completely attenuated in RPMI 8226 although synergistic enhancement of apoptosis was observed in NCI-H 929. Furthermore, following isolation of PKH26^Hi^ quiescent cell populations, TRAIL responses and enhancement of TRAIL responses by GSK343 were completely attenuated. These studies show that EZH2 inhibition enhances TRAIL responses both in TRAIL-sensitive and TRAIL-resistant MM cell lines suspension culture and also in 3D cell culture to model the semi-solid Multiple Myeloma lesions in bone. Pre-existing TRAIL resistance was also enhanced by EZH2, and although synergistic enhancement of TRAIL responses by GSK343 was seen in NCI-H 929, responses were completely lost in TRAIL-resistant RPMI 8226 and also in quiescent cells. These studies highlight that although EZH2 inhibitors enhance TRAIL responses, acquired TRAIL resistance, and the presence of quiescent cells may mediate TRAIL-insensitivity in response to GSK343.

## Introduction

Multiple myeloma is a plasma cell haematological malignancy characterised by neoplastic growth in the bone marrow and is the final stage in a multi-step progressive disorder representing 1% of all cancers and 10% of haematological malignancies (Rajkumar, 2020). Myeloma is characterised by multiple genetic changes, including hyperdiploidy, the non-random gain of a specific subset of chromosomes, translocation of the *IGH* gene leading to aberrant expression of translocated oncogene, and loss and gain of specific chromosomal regions (Morgan, Walker & Davies, 2012). A subset of these myeloma-associated translocations involves the histone methyltransferase MMSET typically t(4;14), and subsequent secondary genetic events are linked to disease progression (Morgan, Walker & Davies, 2012). There is no cure for Multiple Myeloma and current therapies rely on Autologous Stem Cell Transplant and combination treatments with chemotherapeutic agents mainly targeting the intrinsic apoptotic pathway (Rajkumar & Kumar, 2020). Treatment-refractory tumour cells remain a continuous problem in Multiple myeloma therapy with treatment resistant and dormant myeloma cells being implicated in disease relapse, making development of a treatment targeting these cells vital.

TRAIL (Tumour necrosis factor related apoptosis inducing ligand) is an apoptosis-inducing ligand expressed on immune cells, stimulating death receptors 4 and 5 (DR4 and DR5) via the extrinsic apoptosis pathway by DISC (death-inducing signalling complex) induction (Pan *et al*, 1997). This apoptosis pathway is triggered independently of p53 which is commonly mutated in many drug-resistant cancers (Herbst *et al*, 2010). TRAIL also triggers the intrinsic apoptosis pathway and exerts effects on malignant cells with little off target effects on non-cancerous cells making TRAIL mimetics a promising therapy for many cancers. In Multiple Myeloma, DR4 and DR5 are overexpressed (Jourdan *et al*, 2009) making myeloma a good candidate for TRAIL treatment. However, expression of decoy receptors DcR2 and OPG in Multiple Myeloma inhibits the action of TRAIL mimetics (Shipman & Croucher, 2003), as well as c-Flip overexpression, therefore TRAIL sensitisers are being considered for combination therapy (Sayers and Cross, 2014).

EZH2 (Enhancer of Zest Homologue 2) is a histone methyl transferase (HMT) and the catalytic domain of the PCR2 (Polycomb Repressor 2 Complex). EZH2 is responsible for methylation of histone N-terminal tails at site H3K27 (Histone 3 Lysine 27) leading to transcriptional repression. EZH2 is found to be overexpressed in Multiple Myeloma (Croonquist and Van Ness, 2005; Kalushkova *et al*, 2010) as well as many other cancers and has also been found to be involved in cancer progression (Drelon **et al*,* 2016; Verma et al, 2012), acting by repressing the transcription of tumour suppressor genes e.g. TGFBR2 in non-small cell lung carcinoma (Lo Sardo et al, 2021) and LATS2 in Acute myeloid leukaemia (Feng, Hu, Li, Peng & Chen, 2020) and through silencing of pro-apoptotic miRNAs e.g. miR-205 and miR-31 (Zhang et al, 2014). EZH2 overexpression is also linked with a worse prognosis in patients with Multiple Myeloma (Pawlyn *et al*, 2017). GSK343 is a potent, highly selective EZH2 inhibitor, inhibiting epigenetic repression of transcription (Verma *et al*, 2012). EZH2’s action in Multiple Myeloma is thought to be through its dysfunctional interaction with the MMSET protein, both of which are aberrantly overexpressed in Multiple Myeloma due to the t(4;14) translocation placing the MMSET gene downstream of the strong IgH promotor (Popovic **et al*,* 2014). MMSET and EZH2 epigenetic dysfunction have been linked to cancer progression through the impaired regulation of the expression of oncogenes and tumour suppressor genes. Inhibiting EZH2 has proven to be a promising therapeutic with Tazamtostat (EZH2 inhibitor) being recently approved for clinical use in epithelioid sarcoma (Hoy 2020). GSK343 in combination with TRAIL has been found to target drug resistant cancer stem cells by upregulating DR4/5 in colon cancer (Singh *et al*, 2021). EZH2 inhibition successfully induces apoptosis in human Multiple Myeloma cells (Kalushkova *et al*, 2010) and furthermore, distinct metabolic signatures are associated with EZH2 sensitivity (Nylund et al, 2021). Given that some Multiple Myeloma cells have MMSET translocation, and that EZH2 inhibition has been shown to be pro-apoptotic in myeloma cells, the present study set out to determine the potency of GSK343 in sensitising cells to TRAIL.

Previously, we established that Histone deacetylase inhibitor, SAHA potently sensitized TRAIL resistant and sensitive myeloma cells to TRAIL induced apoptosis (Arhoma **et al*,* 2017) and showed that in this 3D cell culture model to mimic the semi-solid lesions observed in bone, SAHA enhanced TRAIL sensitisation. We have extended these studies to GSK343 in the current study. Since TRAIL resistance emerges rapidly in cancers, it is also important to evaluate the ability of GSK343 to sensitise TRAIL resistant cells to TRAIL induced apoptosis. TRAIL resistance emerges through a variety of mechanisms including constitutively active Akt and overexpression of c-FLIP (Sayers and Cross, 2014). Prolonged exposure to TRAIL has been found to lead to TRAIL resistant CD138 negative cells, and TRAIL in combination with Doxorubicin has been found to successfully eliminate these cells (Vitovski *et al*, 2012). These quiescent cells are also likely responsible for resistance to current therapies used to clinically treat Multiple Myeloma such as bortezomib, whereby bortezomib induces quiescence through the unfolded protein response by attenuating eIF2alpha phosphorylation preventing pro-apoptotic gene induction (GADD153) and upregulation of GRP78 (pro-survival chaperone) (Adomako *et al*, 2015; Schewe & Aguirre-Ghiso, 2009). Therefore, the present study aimed to determine if GSK343 in combination with TRAIL would induce apoptosis, whether enhanced effects extended to cells with acquired TRAIL-resistance, and whether this approach might target quiescent cells that are known to mediate treatment resistant in Multiple Myeloma cells.

## Materials and methods

### Cell lines and cell culture

Five established Human Multiple Myeloma cell lines (RPMI 8226, OPM2, NCI-H 929, U266 and JJN3) and one cell line (ADC-1) generated from a plasma cell leukaemia were used: RPMI 8226 (ECACC (European Collection of Cell Cultures) Salisbury, UK, cat no: 87012702) NCI-H 929 (EACC, Salisbury, UK Cat no. 95050415), OPM2 (Deutsche Sammlung von Mikroorganismen und Zellkulturen GmbH (DSMZ), Braunschweig, Germany; DSMZ no ACC 50. U266 (purchased from LGC Standards (UK)), JJN3 plasma cell leukaemia cells were gifted by Professor I Franklin (University of Glasgow). ADC-1 from the peripheral blood of a plasma cell leukaemia patient with suitable ethical consent approved by the South Sheffield Research Ethics committee (REC reference: 05/Q2305/96) (Department of Haematology, Sheffield Teaching Hospital, UK).

All Multiple Myeloma cell lines were cultured under standard culture conditions; 37°C with 5% CO_2_ atmosphere in RPMI-1640 medium + L-Glutamine and supplemented with supplemented with 1% penicillin–streptomycin, 1% non-essential amino acid, and 10% foetal calf serum. Cells were tested for mycoplasma infections throughout the study (MycoAlertTM Mycoplasma detection assay).

Multiple Myeloma cells were seeded into complete RPMI-1640 medium and maintained in a T75cm2 flask (Invitrogen). All cells were incubated under standard cell culture conditions and seeded biweekly at 1:5 split ratio split. For seeding in 96 well plate cells were centrifuged at 400g for 5 minutes, resuspended in complete media and counted using trypan blue exclusion using a countless system (Invitrogen) 200μl of cells at 350,000 cells/ml was transferred into a 96 well plate and incubated for 2-3 hours before addition of therapeutic agents.

### Treatment with TRAIL and GSK343

For combinations studies with GSK343, cell lines were treated concentrations of TRAIL that did not significantly induce apoptosis based on our previous study (Arhoma *et al*, 2017) as follows: OPM-2 (0.5ng/ml), RPMI 8226 (2ng/ml), NCI-H 929 (2ng/ml), ADC-1 (2ng/ml), U266 (50ng/ml), and JJN3 (50ng/ml). GSK343 was initially diluted to 10mM in DMSO and further diluted in complete medium. Cells were treated with 0-20μM and control cells treated with 0.5% v/v DMSO vehicle control. Cells were treated for 24 hours with either vehicle control, TRAIL (at the specified dose), GSK343 (0-20 μM) or combined treatment.

### Assessment of apoptosis by nuclear morphology

After treatment with GSK343 +/− TRAIL for 24 hours apoptosis induction was assessed by Hoechst 33343 and propidium Iodide (PI) stain (Sigma-Aldrich, Dorset, England). Cells were incubated with 10μg/ml Hoechst 33342 and 10μg/ml PI at 37°C for 30 minutes examined using IX81 Olympus fluorescence microscope, using Cell F software for image capture. Images were captured using a UC30 colour camera and triple filter for DAPI/FITC/TRITC. Images containing at least 100 cells were obtained from duplicate wells in three technical repeats and apoptosis calculated by manual counting of cells with characteristic condensed or pyknotic nuclei. PI was used to exclude necrosis.

### Assessment of ATP as a measure of cytotoxicity

To measure ATP levels and so the relative cell viability after treatment with TRAIL+/− GSK343, cells (350,000 cells/ml) were transferred into a 96 well plate and treated with GSK343 +/− TRAIL for 24 hours, each treatment group containing three replicates with three independent repeat experiments. 50μl CellTiter-Glo^®^ Reagent was then added to each well and incubated at room temperature for 10 minutes. Luminescence was then measured using Wallac Victor 2 1420 luminometer.

### Assessment of Caspase-3 activity

The NucView Caspase-3 activity assay (Biotium, Cambridge Biosciences, Cambridge, UK) was used to confirm apoptosis observed by Hoechst 33423/PI staining in selected cell lines (RPMI 8226 and ADC-1). Negative control cells were produced by treatment with 10 μM Z-DEVD-FMK caspase-3 inhibitor (R&D systems). 200μl of cell suspension from each cell line including the negative control group was transferred into a flow cytometry tube before being treated with 2.5μl of 0.2mM NucViewTM 488 caspase-3 substrate (fluorogenic DNA dye and DEVD moiety substrate for caspase-3) to allow detection of caspase-3 and visualisation of nuclear morphology. These cells were then incubated at room temperature for 20 minutes and analysed on Beckman Coulter Gallios flow cytometer acquiring ten thousand events per sample.

### Assessment of Caspase-8 and Caspase-9 activity

Caspase-8 and caspase-9 activities were assessed in all 6 Multiple Myeloma cell lines. Cells were treated with the pre-determined dose of TRAIL in the presence or absence of GSK343 (0, 5, 10 and 15μM) in triplicate wells of in white, Fisher Scientific 96 well plates. Cells were lysed as the Caspase-Glo^®^ reagent was added the assay was performed following the manufacturer’s instructions with 24-hour incubation in Caspase-Glo reagent. 50μl of either caspase-8 reagent or caspase-9 (Promega) reagent was then added with MG132 proteosome inhibitor. The samples were then incubated at room temperature for 90 minutes before the Wallac Victor 2 1420 luminometer was used to measure the luminescence.

### Generation of TRAIL resistant cell lines

NCI-H 929 and RPMI 8226 TRAIL-sensitive cells were treated a period of 12 months with increasing concentrations from 2ng/ml to 50ng/ml of TRAIL, being increased depending on weekly cell viability to produce TRAIL resistant cell lines. Live cells were counted using Invitrogen’s Cell Countless system and viable cells were determined using trypan blue staining. Cells were then treated with GSK343 +/− TRAIL at 50ng/ml and assessed for apoptosis using Hoechst 3334/PI stain.

### Assessment of the effect of TRAIL +/− GSK343 using 3D cell culture alginate sphere assay

Cells (1×10^6^ cells/ml) were resuspended in 1.2% medium viscosity sodium alginate dissolved in saline (sigma). The cells in alginate solution were dropped into 200mM CaCl_2_ through a 19-gauge needle and allowed to polymerise for 15 minutes at 37°C. Beads were then washed three times; twice with NaCl (0.15M) and once in the media. The beads were then left in the media in the incubator (37°C) to form spheroids. The medium was changed twice a week and the spheroids maintained for 10-14 days. After this point the spheroids were treated with the pre-determined dose of TRAIL +/− GSK343 depending on the cell line for 24 hours and then stained with Hoechst 33342/PI. Each 1ml of cell suspension generated approximately 40 polymerised beads, each bead containing approximately 2.5×10^4^ cells.

Alginate spheroids were imaged every 2-3 days using an Olympus IX81 fluorescence microscope and a Hoechst 33342/PI staining to determine cell viability, morphological analysis, and diameter of the spheroids. To image the spheroids the alginate beads were first dissolved and incubated in alginate dissolving buffer containing 30mM EDITA (Fisher), 55mM sodium citrate (sigma) and 0.15M NaCl (pH6) for 15 minutes followed by centrifugation for 5mins (600g) and tumour spheres resuspended in media and imaged as described previously with Hoechst 33342/PI.

### Assessment of TRAIL +/− GSK343 in quiescent cells: PKH26 staining

The PKH26 membrane labelling kit (Sigma) was used to stain the Multiple Myeloma cells and subsequently isolate quiescent cells. The cells were first centrifuged at 400g for 5 minutes and resuspended in medium without FCS, centrifugation was repeated and supernatant aspirated and 125 μl of dilutant C from the kit was added. 5 μl of PKH26 dye solution was added (30 μl ethanol and 4 μl 1mM PKH26 linker) and the samples incubated for 1 minute at room temperature before the addition of 130 μl FCS to stop the reaction followed by a 10-minute centrifugation. The cells were then washed twice in 10ml of media and re-suspended in 2ml media and seeded into 6 well plates. The media was changed 3 times a week.

Assessment of PKH26 stained cells after 10-14 days incubation assessment for stain intensity was performed using an Olympus IX81 fluorescent microscope. Images were captured using Olympus Cell-F software taking a dual image of brightfield and Texas Red filters and further analysed by flow cytometry (BD FACSCaliburTM, BD Biosciences). Day 0 samples (stained and unstained) were used for baseline measurements for the flow cytometry software. Flow cytometry measured the cell granularity (side scatter) and the cell size (forward scatter), identifying the viable cell population. A histogram plot comparing the percentage of stained to unstained cells of a gated population. Ten thousand events acquired per sample; samples from each line were assessed three times a week, data analysis performed using FlowJo software. 14 days after staining, the PKH26 labelled Multiple Myeloma cell lines were sorted into PKH26^Hi^ and PKH26^Lo^ populations, gating around the unstained population using an unlabelled cell sample and PKH26^Hi^ were classed as having stain levels comparable to day 1 using a Becton Dickinson FACSAria at The University of Sheffield’s flow cytometry facility.

### Assessment of apoptosis in PKH26^Hi^ cells in response to TRAIL +/− GSK343

GSK343 induced TRAIL response was measured in quiescent cells using Annexin-V-FITC/Hoechst 33342 staining to measure apoptosis and assessment of both nuclear morphology and further assessment of Annexin-V-FITC positivity. Initial preliminary studies showed an absence of response at low doses of TRAIL (2ng/ml) so all subsequent experiments used 50ng/ml, the maximum dose used for TRAIL-resistant cell lines. After treatment for 24 hours, cells were centrifuged at 400g for 5 minutes, supernatant removed and 100 μl Annexin V binding buffer (10 mM HEPES/NaOH, 140 mM NaCl, 2.5 mM CaCl_2_ at pH 7.4) added. 10 μl Hoechst 33342 and 5 μl Annexin V-FITC stain were then added and left to incubate for 20 minutes. Images were captured on a UC-30 camera on an Olympus IX-81 microscope using a triple filter and captured as described previously. Since The strong PKH26 stain in PKH26^Hi^ cells partially obscured the Hoechst 33342 and Annexin-V-FITC signal so additional imaging using a DAPI filter was performed to ensure apoptosis was not under-estimated in the PKH26Hi population.

### Statistical analysis

For all experiments, a median with range was calculated and Shaprio-Wilke test was used (Stats Direct Ltd, England) to determine if the data followed a normal distribution. Kruskal-Wallis and Connover-Inman post hoc were used to determine whether the data not falling within a normal range was of a significant difference (p<0.05). All data represent combined data from 3 technical repeats with the exception of quiescence cell data (fig 6c-d) where data shown is from a representative analysed experiment (n=3 wells) from just 2 technical repeats.

## Results

### GSK343 enhances TRAIL responses in Multiple Myeloma cell lines

Multiple Myeloma cell lines were treated with increasing concentration of GSK343 (0, 1, 5, 10, 15 and 20μM) in combination with and without a previously determined dose of TRAIL known to induce only low levels of apoptosis (Arhoma *et al*, 2017) and apoptosis determined by Hoechst 33342/ PI stain. GSK343 alone induced apoptosis in NCI-H 929, RPMI 8226 and ADC-1 cells in the higher doses (15-20 μM). U266 cells were particularly sensitive to GSK343 alone with apoptosis being significantly induced (≥10μM GSK343) (fig 1a). JJN3 and OPM2 were not sensitive to GSK343 without TRAIL. All cell lines when exposed to TRAIL and GSK343 experienced a synergistic response of induced apoptosis, with the apoptotic effect observed differing depending on the cell line and dose of GSK343, e.g., in RPMI 8226 cells, a significant difference in apoptosis was not observed in the combination treatment when compared to GSK343 alone below 10μM GSK343, after which a significant synergistic response was observed. In JJN3 cells however a significant synergistic response was observed with just 1μM GSK343. Synergistic enhancement of apoptosis was observed only at 5μM GSK343, beyond which GSK343 alone resulted in >90% cell death and loss of synergistic responses. TRAIL-resistant OPM-2 showed weak but significant induction only at 10μM GSK343.

**Figure 1:**
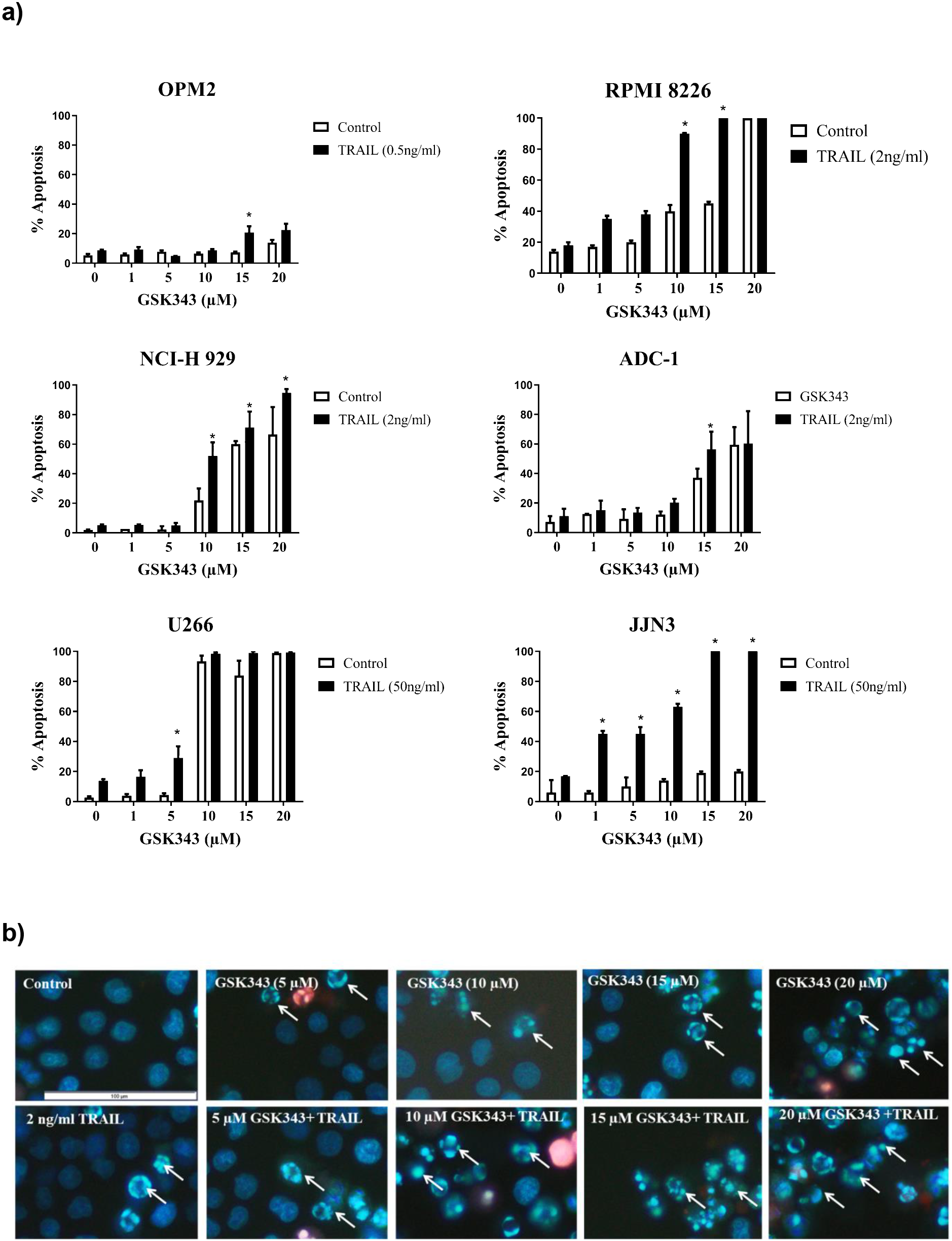
Apoptotic response to the synergy of GSK343 0-20μM +/− TRAIL measured by Hoechst/PI stain. a) Morphological assessment of apoptosis (Hoechst 33342/PI stain) showing percentage apoptosis in response to drug treatment with varying GSK343 concentrations. The synergistic response was defined as a combination of GSK343, and TRAIL compared to the sum of TRAIL and GSK343 alone with significance determined by Kruskal-Wallis test (p< 0.05) with significantly higher levels of apoptosis induced by the synergistic response to GSK343 and TRAIL. b) Representative RPMI 8226 cell morphology analysis using Hoechst 33342/PI stain presenting the synergistic effect of GSK343 and TRAIL in inducing apoptosis.

### The effect of GSK343 +/− TRAIL ATP levels as a marker of cytotoxicity

The response of the Multiple Myeloma cell lines to GSK343 and TRAIL and GSK343 0-20μM showed a dose-dependent significant decrease in ATP levels (fig 2) consistent with cytotoxicity. The synergistic effect of TRAIL and GSK343 had differing effects on the ATP levels of the different cell lines. The treatment with GSK343 and TRAIL induced a reduction of ATP levels in cell lines: OPM2, RPMI 8226, NCI-H 929 and U266. Whereas cell lines JJN3 and ADC-1 were most effected by GSK343 alone with a significant decrease in ATP levels, however no synergistic response was observed when combined with TRAIL.

**Figure 2:**
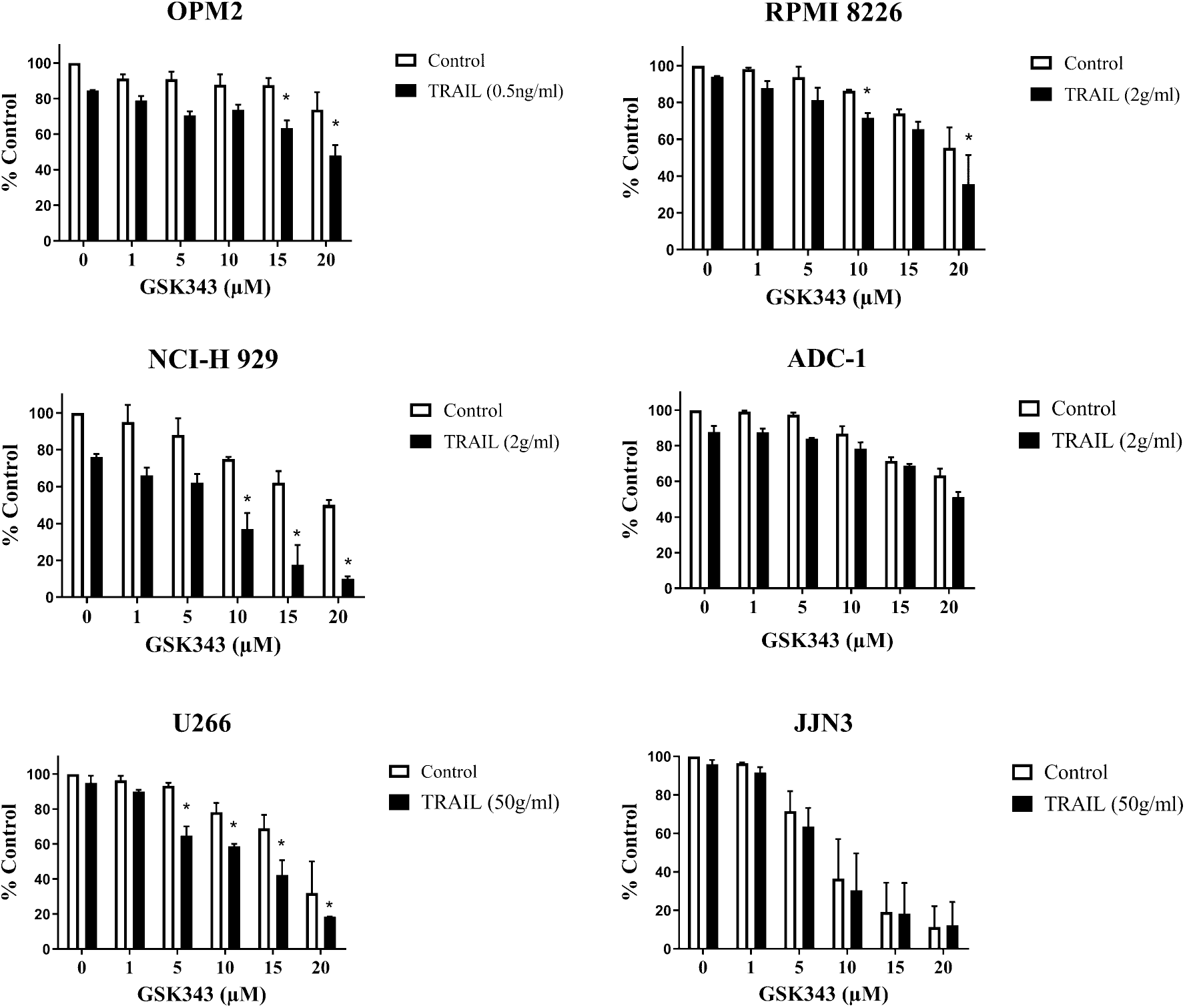
Cell viability analysis of Multiple Myeloma cell lines measuring ATP levels, as evaluated by the CellTiter-Glo assay. The data was compared to a vehicle control representing 100% viability and expressed as a median with range of the three experiments in triplicate. Significant enhancement of TRAIL responses by GSK343 is represented by an asterisk (*). Statistical significance determined by Kruskal-Wallis and set at p<0.05.

### Caspase activity in Multiple Myeloma cell lines in response to GSK343 treatment +/− TRAIL

To confirm apoptosis in response to treatments, the NucView 488 capsase-3 activation assay was used to determine caspase-3 activity and analysed by flow cytometry (figure 3a). NCI-H 929, RPMI 8226 and U266 showed particularly potent caspase-8 activity induction although much of this was in response to TRAIL alone consistent with death-receptor mediated apoptosis. GSK343 alone did not induce potent caspase-8 activation although significant induction was observed in U266, JJN3 and OPM2. Potent induction of caspase-9 activity was observed in U266 and RPMI 8226 in response to combination treatment.

**Figure 3:**
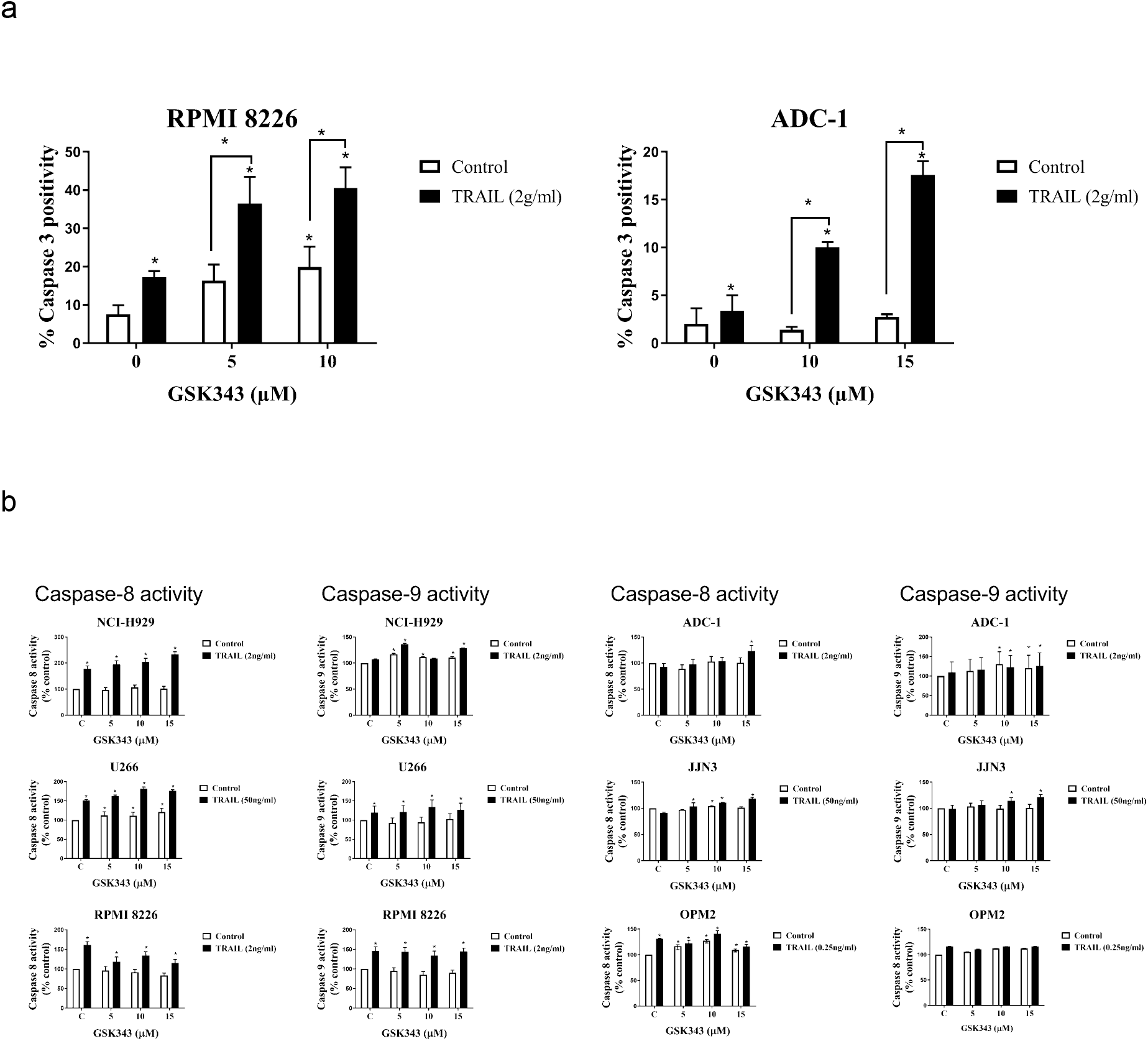
The effect of GSK343+/− TRAIL on caspase activity in Multiple Myeloma cell lines. a) Caspase 3 levels in ADC-1 and RPMI 8226 cell lines after GSK343 and TRAIL alone and combined treatment for 24hrs GSK343 (0-15μM). b) Caspase-8 and −9 activity after treatment with increasing concentrations of GSK343 +/−TRAIL from three independent experiments performed in triplicate using Caspase-Glo^®^ assay in different Multiple Myeloma cell lines compared with a vehicle control (100%). All data expressed as a median with range and significance determined using Kruskal-Wallis p< 0.05 by comparison to the vehicle control.

### Effects of TRAIL +/− GSK343 3D cell culture

3D cell culture models of Multiple Myeloma cell lines were generated using alginate bead culture as described previously (Arhoma *et al*, 2017). U266 (TRAIL-resistant) and NCI-H 929 (TRAIL sensitive) were chosen for 3D cell culture studies. 3D cell cultures of both NCI-H 929 and U266 were treated with GSK343 alone and in combination with TRAIL for 24 hours and apoptosis measured using Hoechst 33342/PI stain (figure 4a-b). U266 was treated with GSK343 (5 μM) +/− 50ng/ml TRAIL as this dose showed synergistic enhancement of this cell line in suspension culture, similarly GSK343 (10 μM) +/− 2ng/ml TRAIL was used for NCI-H 929. In both cell lines, both TRAIL responses and GSK343 responses were significantly increased in 3D cell culture *vs.* suspension culture, and in U266 TRAIL responses were potently and synergistically enhanced in this TRAIL-resistant cell line in 3D cell culture (Fig 4a)

**Figure 4:**
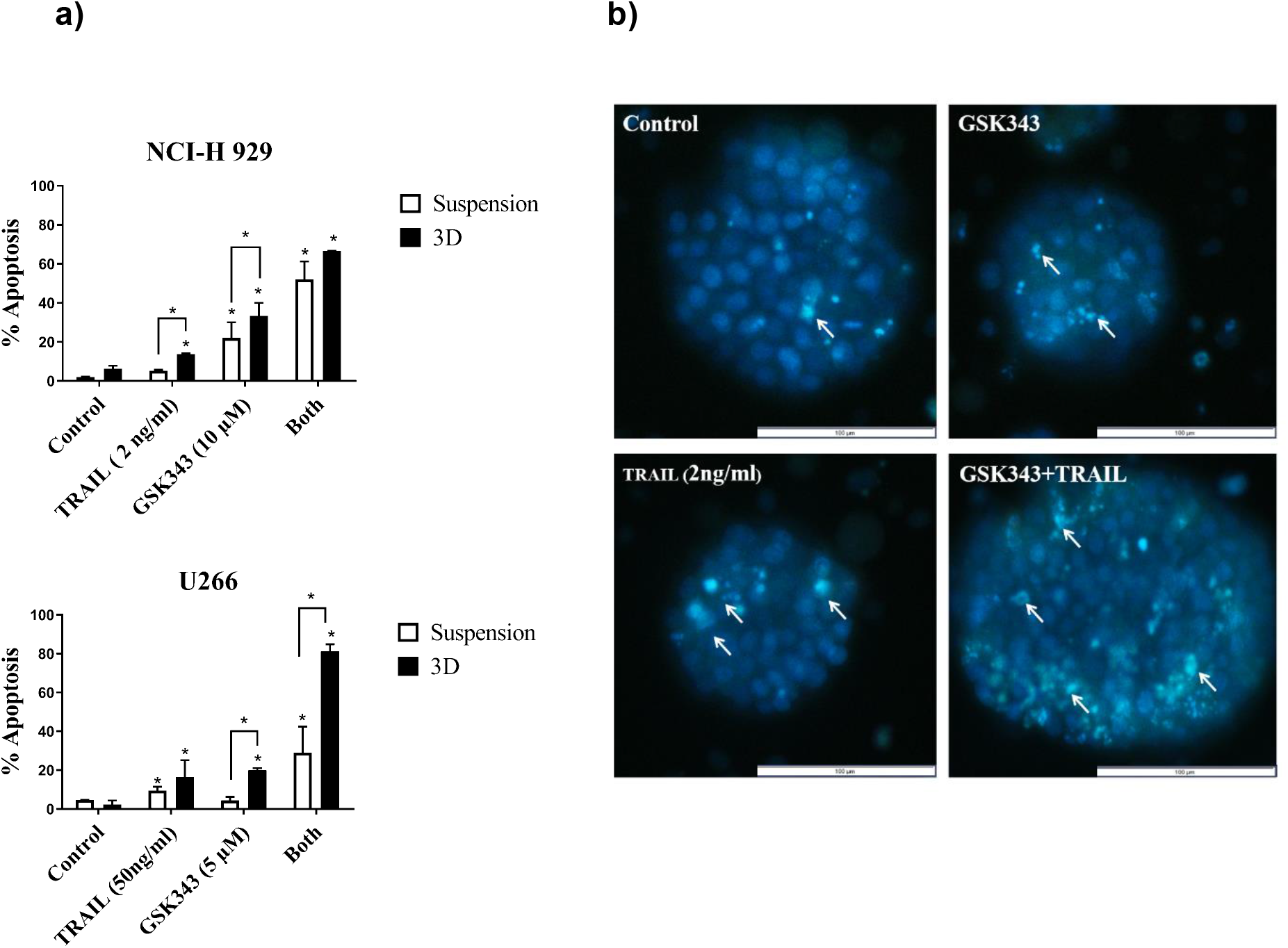
The effect of GSK343, TRAIL and combination treatment on apoptosis in suspension culture vs. 3D cell culture in NCI-H 929 and U266 measured by Hoechst 33342/PI staining. Data expressed as a median with range fromtriplicate results fromthree experiments statistical significance in comparison to control was defined as p< 0.05. b) Morphological assessment of NCI-H 929 in representative spheroids using Hoechst 33342/PI stain to measure apoptosis with GSK343 +/−TRAIL. Arrows indicate selected apoptotic cells with condensed nuclei.

### Effects of TRAIL +/− GSK343 in TRAIL-resistant cell populations

The generation of TRAIL-resistant cell populations was performed in TRAIL-sensitive NCI-H 929 and RPMI 8226 through prolonged exposure to TRAIL of increasing concentrations over the course of 12 months. The cell viability analysis (Fig 5a) and apoptosis assessment (Fig 5b) confirmed the decreased TRAIL responses in both cell lines. When treated with TRAIL at 50ng/ml, a dose that would induce apoptosis in the majority of cells, neither cell line showed significant apoptosis in the TRAIL-resistant populations. Combination treatment TRAIL and 10μM GSK343 resulted in significant attenuation of GSK343 responses in both cell lines, and potent inhibition of apoptosis in RPMI 8226 in response to combination treatment. Responses to combination treatment were attenuated in NCI-H 929 in TRAIL-resistant cells although synergistic enhancement of TRAIL responses by GSK343 was still present with apoptosis levels similar to parental cells after dual treatment (Fig 5c).

**Figure 5:**
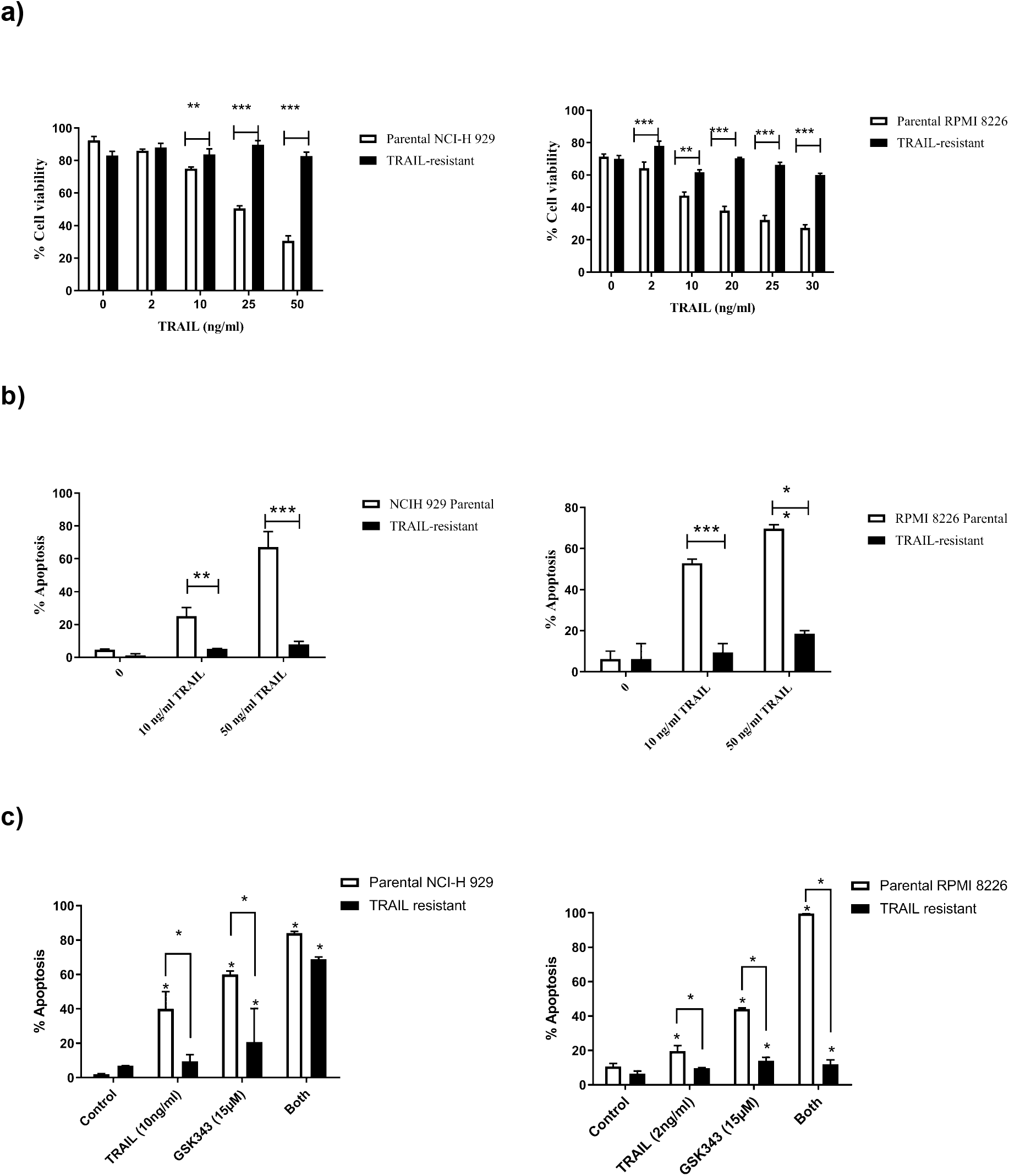
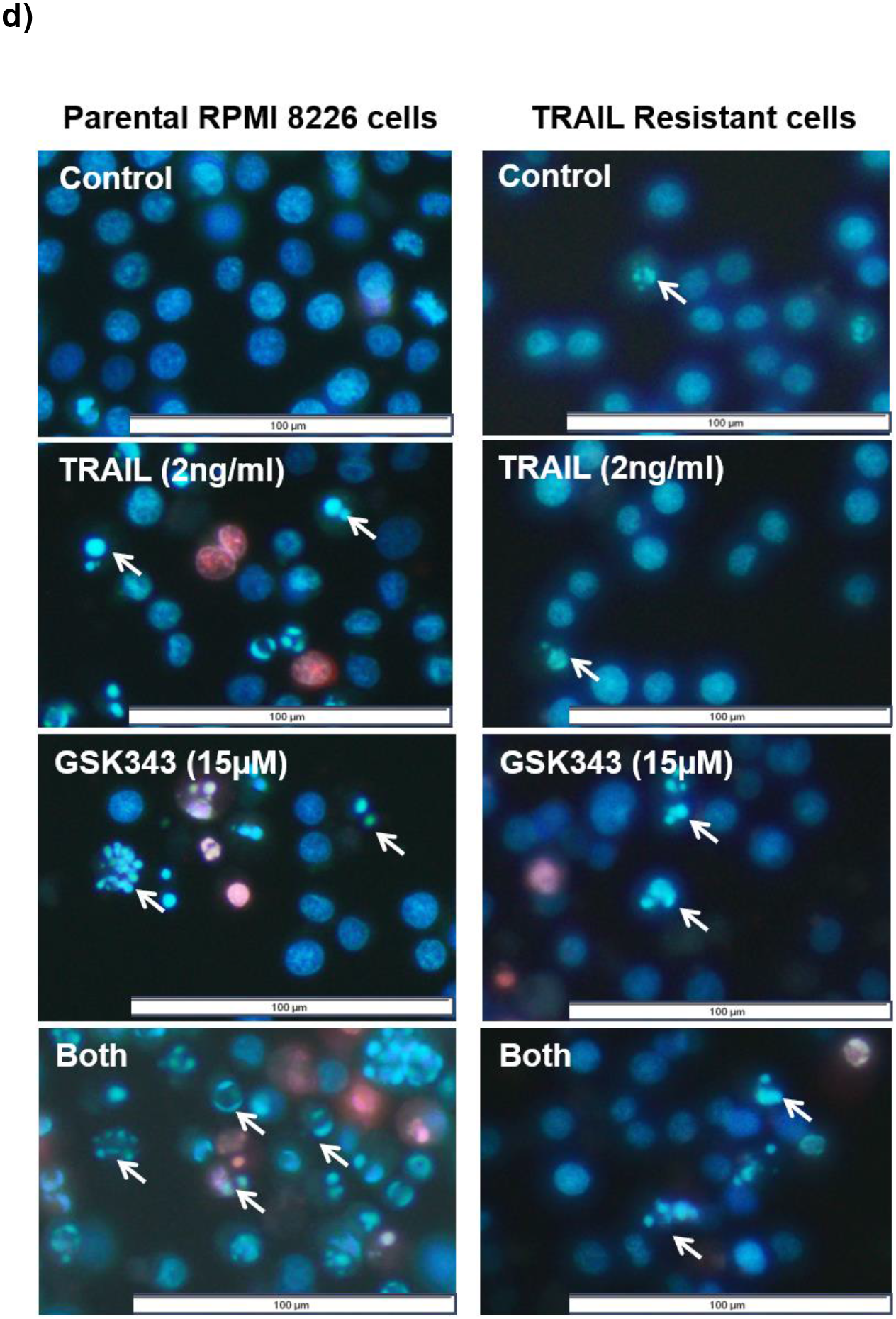
a) Chronic exposure of increasing TRAIL concentration over 12 months resulted in TRAIL-resistant populations showing increased cytotoxicity as determined by ATP levels and b) decreased apoptosis in response to TRAIL in the previously TRAIL-sensitive NCI-H 929 and RPMI 8226 as determined by Hoechst 33342/PI staining. c) TRAIL-resistant cells show reduced GSK343 sensitivity and reduced sensitivity to combination treatment, although synergistic enhancement of TRAIL by GSK343 still occurred in NCI-H 929. d) Representative treatment of parental and TRAIL-resistant RPMI 8226 with TRAIL +/− GSK343. Statistical significance determined by Kruskal-Wallis test: (*= p< 0.05, **= p< 0.01 and ***= p< 0.001).

### Assessment of TRAIL +/− GSK343 responses in PKH26^Hi^ quiescent Multiple Myeloma cells

Since PKH26 dye is lost from cells by cell division, and quiescent cells are non-proliferative, Multiple Myeloma cells were stained with PKH26 and PKH26HI populations isolated after 14 days. NCI-H 929, RPMI 8226 and U266 cell lines were used to isolate PKH26Hi non-proliferative cell populations. All cell lines by day 14 had a minority of cells that were strongly positive for PKH26 (Fig 6a) as determined by flow cytometry. PHK26Hi and PKH26Lo cells were isolated by Flow Activated Cell Sorting (FACS) as shown in Fig 6b, showing successful sorting of populations with approximately 98% purity. Isolated PHK26Hi and PKH26Lo cells were then treated with TRAIL +/− GSK343 and apoptosis assessed, and results compared to unsorted cells. Initial experiments from the TRAIL-sensitive NCI-H 929 and RPMI 8226 showed a lack of response in PKH26Hi cells, so cells were subsequently treated with 50ng/ml TRAIL, the dose used on TRAIL-resistant cell populations. Figure 6c-d shows that in the presence of TRAIL (50ng/ml) and GSK343 (20nM), unsorted cells were efficiently killed at this high dose of TRAIL, GSK343 (20nM) also killed the majority of cells and combination treatment resulted in approximately 80% cell death in 24 hours (n=2), In contrast, PKH26Hi cells were resistant to TRAIL, GSK343 and combination treatment (Fig 6d).

**Figure 6:**
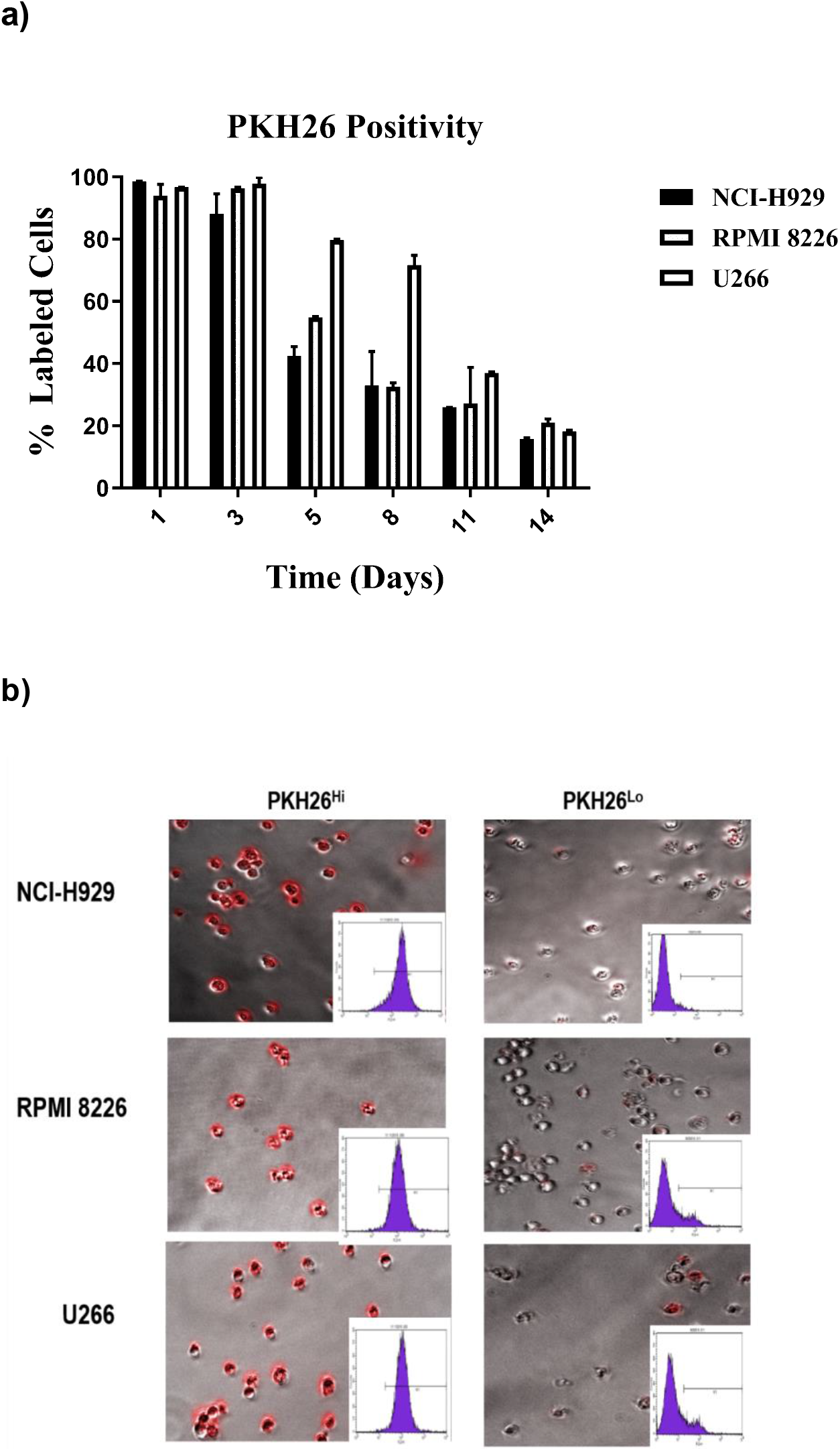
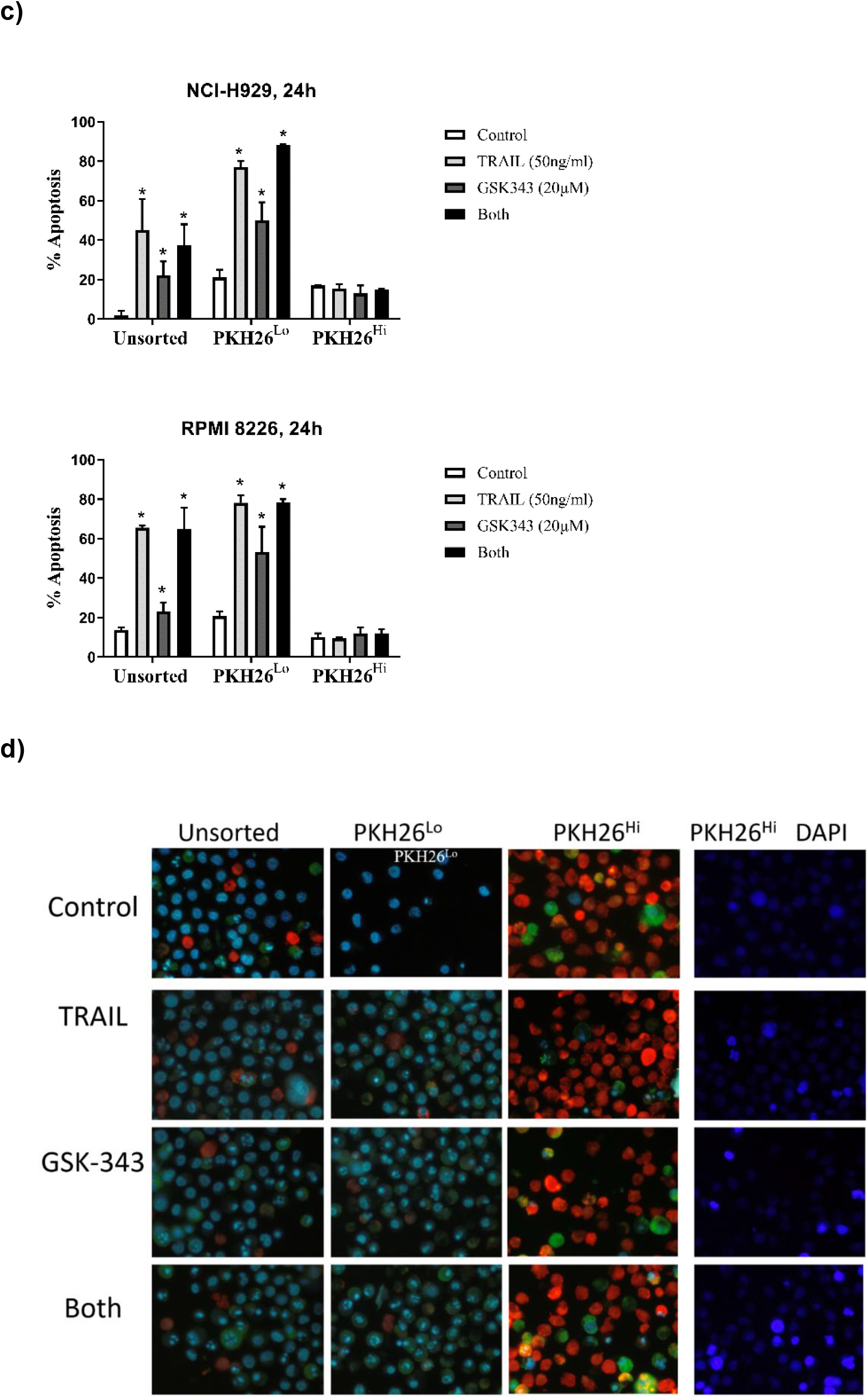
PKH26 fluorescent positivity in myeloma cells determined by flow cytometry and FACS a) Percent positivity of NCI-H 929, RPMI 8226 and U266 showing decrease in PKH26 positivity as determined by flow cytometry. b) Post-sorting of 14-day stained Multiple Myeloma cells showing purity of populations, c) responses of unsorted, PKH26 ^Lo^ and PKH26^Hi^ cells in response to TRAIL (50ng/ml) and GSK343 (20ng/ml) and d) representative images of unsorted, PKH26^Lo^ and PKH26^Hi^ cells treated with TRAIL (50ng/ml) and GSK343 (20ng/ml) as determined by Hoechst 33342 and Annexin-V-FITC staining. For improved visualisation of nuclear morphology, nuclear morphology of PKH26Hi cells are also shown for the DAPI channel only.

## Discussion

Despite the introduction of many novel therapies for Multiple Myeloma, refractory disease eventually develops, and Multiple Myeloma remains uncurable for the majority of patients. Clinical trials using modified TRAIL in combination with thalidomide shows an acceptable safety profile and some promising activity (Geng et al, 2014). Furthermore, combination of modified TRAIL with Dexamethasone shows increased progression free survival in refractory disease (Leng et al, 2017). Furthermore, dual inhibition of DNA methyltransferases and EZH2 can overcome resistance of Thalidomide derivatives in Multiple Myeloma, highlighting the importance of EZH2 targeting in Multiple Myeloma (Dimopoulus et al, 2018). Support for this comes from a clinical trial of EZH2 inhibitor GSK2816126 in haematological malignancies including Multiple Myeloma. The trial resulted in GSK2816126 presenting a lack of therapeutic potential at the maximum allowed dosage (Yap et al, 2019). Because of the potent effects of EZH2 inhibition observed in vitro, this suggests that clinically, combination treatment may be more effective.

TRAIL responses can be enhanced by a wide range of potential anti-tumour agents including Histone deacetylase inhibitors, proteasome inhibitors, nuclear export inhibitors, Bh-3 mimetics and (Cross and Sayers, 2014, Phillips et al, 2019). We have previously shown that HDAC inhibitors can sensitise Multiple Myeloma cells to TRAIL (Arhoma et al, 2017) whilst Braun et al (2015) showed increased TRAIL signaling in lymphomas using a pan-histone methyltransferase. EZH2 inhibitor UNC1999 was found to reduce cell viability as well as reduce the expression of oncogenes: XBP-1, PRDM1 and c-MYC in Multiple Myeloma cell lines (Alzrigat **et al*,* 2017). This is also supported by another EZH2 inhibitor, DZNep being found to induce apoptosis in human multiple myeloma cells when used as a single agent (Kalushkova *et al*, 2010). In this study we show for the first time that the EZH2-specific histone methyltransferase inhibitor GSK343 enhances TRAIL responses in Multiple Myeloma cells and those responses extend to 3D cell cultures, mirroring recent findings using HDACi (Arhoma et al, 2017). Furthermore, since TRAIL is known to result in the rapid emergence of TRAIL-insensitive populations, we assessed two known mechanisms of TRAIL-insensitivity: TRAIL-resistance induced by TRAIL treatment, and assessment of quiescent cells which are known to be TRAIL-resistant in Multiple Myeloma (Vitovski et al, 2012)

The EZH2 inhibitor GSK343 in combination with TRAIL synergistically or additively induced apoptosis in all cell lines, and U266 was particularly sensitive to GSK343. EZH2 is overexpressed in many solid tumours (Varambally et al, 2002) and haematological malignancies, including Multiple Myeloma (Croonquist and Van Ness, 2005; Kalushkova *et al*, 2010) being correlated with poor prognosis. EZH2 has been identified as a promising therapeutic target for cancer through its interaction with MMSET, with EZH2 repressing and MMSET activating transcription (Asangani *et al*, 2013). This dysfunction is thought to arise through the t(4;14) MMSET translocation in Multiple Myeloma, placing it under the control of a strong IgH promotor (Popovic *et al*, 2014). EZH2 inhibition has previously been found to reduce c-MYC expression (Popovic *et al*, 2014) and so, it was hypothesised that cells with the MMSET translocation (OPM2, NCI-H 929 and ADC-1) would be more responsive to GSK343 treatment than MMSET negative RPMI 8226, U226 and JJN3 cells. However, the results from the present study do not support this hypothesis, with no correlation between MMSET and either GSK343 sensitivity, or GSK343-enhancement of TRAIL responses (fig 1–2). These results suggest that the interplay of EZH2’s epigenetic activity in Multiple Myeloma may be more complex than its interaction with MMSET alone and suggestive of a more generalised disruption to normal epigenetic programming in Multiple Myeloma, highlighting EZH2 as a valid target for Multiple Myeloma. Histone methyltransferase inhibition has been found to induce TRAIL responses: Benoit et al (2013) and Braun et al (2015) have found that inhibition of EZH2 via the pan-HMT inhibitor DZNEP enhances TRAIL response through degradation of c-FLIP (Braun *et al*, 2015) as well as more recently Singh et al (2021) found that EZH2 decrease via salinomycin also enhanced TRAIL response in colon cancer stem cells. This is the first study to assess the direct EZH2 inhibitor GSK343 on TRAIL responses warranting further studies into the impact of EZH2 on TRAIL responses in more disease relevant situations including 3D cell culture, TRAIL resistance models and quiescent CSC-like cells.

Our results suggest that GSK343 with or without TRAIL can induce apoptosis, however both TRAIL and GSK343 also induced caspase-9 activation, suggesting activation of the intrinsic pathway but both treatments independently consistent with finding from Argarwal et al (2016) that in RPMI 8226, INA-6 and LP-1 Multiple Myeloma cell lines, EZH2 inhibitors GSK343 and UNC1999 alone stimulate apoptosis via both the intrinsic and extrinsic apoptosis pathway. This suggests that in combination with TRAIL this effect may be amplified through the synergy of the drugs and show potential as Multiple Myeloma treatment as shown in our study.

### GSK343 further enhances TRAIL responses in 3D cell culture

GSK343 alone and in combination with TRAIL was found to induce apoptosis more potently in both NCI-H 929 and RPMI 8226 derived 3D alginate Multiple Myeloma models in comparison to their cell suspension equivalent (fig 4a). Our previous studies have shown enhancement of TRAIL with HDAC inhibitors in both suspension and 3D cell culture with effects potentiated by 3D cell culture (Arhoma **et al*,* 2017). The results of the present study presenting GSK343+/−TRAIL is more potent in inducing apoptosis in 3D cell culture are consistent with findings from epithelial sarcoma cells, where. Hypoxia observed within cell colonies due to the different microenvironments has been implemented in the increase of effect of TRAIL in 3D cell culture through heightened sensitivity of the cell to apoptosis. Hypoxia causes a change in the epigenetic state of cells altering expression of genes encoding molecules controlling cell fate (Batie *et al*, 2019). Hypoxia also stimulates through the expression of HIFs (hypoxia inducible factor family of transcription factors) many pathways that may lead to cell death such as the unfolded protein response (UPR) pathway (Wouters & Koritzinsky *et al*, 2008). Supporting this is findings by Gobbi et al (2010) that hypoxic conditions induce a sensitivity of tumour cells to TRAIL through the down regulation of anti-apoptotic PKCε suggesting that the hypoxic microenvironments within the 3D Multiple Myeloma tumour model were responsible for the observed increased TRAIL sensitivity.

### GSK343-induced TRAIL sensitisation is attenuated in TRAIL-resistant cells

TRAIL resistant cell lines were produced by increasing concentrations of TRAIL over the course of a year. Surviving cells has lost TRAIL sensitivity, showed little response to TRAIL in comparison to parental cells (figure 5a and 5b). GSK343 alone and in combination with TRAIL had little impact on apoptosis in TRAIL resistant cell lines in comparison with the parental RPMI 8226 (figure 5c) although synergistic responses were still seen in NCI-H 929, albeit less potent than in parental cells. This suggests that TRAIL resistance produced from TRAIL exposure induces a more generalised apoptosis resistance, although the mechanism may differ between cell lines. In line with our findings, a previous study found that exposure of Multiple Myeloma cells to TRAIL over a prolonged period of time leads to TRAIL-resistant CD138 negative cells with a reduced response to TRAIL observed (Vitovski *et al*, 2012). Known TRAIL resistance mechanisms are diverse and include c-FLIP overexpression, persistent activation of AKT and modulation of decoy and death receptor expression (Chen **et al*,* 2001; Krueger, Baumann, Krammer & Kirchhoff, 2001, Sayers and Cross, 2014)

### Quiescent cells are insensitive to both TRAIL and GSK343

Quiescent cells are implicated in Multiple Myeloma relapse and resistance. To isolate quiescent cells NCI-H 929 and RMPI-8226 cells were isolated into PKH26Hi and PKH26Lo groups (figure 6) and effect of GSK343+/−TRAIL on apoptosis measured. In all treatment groups, no significant apoptosis was induced in the PKH26Hi cells compared to unsorted and PKH26Lo suggesting that GSK343 alone and in combination with TRAIL has no effect on quiescent cells (figure 6). This suggests that the quiescent PKH26Hi cells presenting resistance to all treatments likely are of the cancer stem cell phenotype may be responsible for the observed treatment resistance supported by Vitovski *et al*’s (2012) findings that CD138 negative cells are more resistant to TRAIL therapy. Apoptotic assessment of PKH26Hi cells was problematic, with PKH26 signal obscuring the Hoechst 33342 signal, hence the use of Annexin-V-FITC to further support findings. We therefore acknowledge that apoptotic assessment might be an under-estimate in PKH26Hi cells. Additionally, the presented data is representative of just two completed experiments. Chen et al (2014) showed that PKH26Hi RPMI 8226 and NCI-H 929 populations presented stem like phenotypes and were resistant to anti-cancer agents such as bortezomib. Quiescent cells such as these, surviving bortezomib treatment have been shown to upregulate GRP78 (transcription factor involved in the unfolded protein response promoting survival) and p21CIP1 whilst downregulation of pRb, CDK6, Ki67 (Adomako *et al*, 2015), implicating the involvement of the unfolded protein response in promoting cell survival in quiescent cancer stem cells. Multiple Myeloma cell which retained DiD, which are analogous with PKH26^Hi^ have been found to be resistant to melphalan (Lawson *et al*, 2015). This suggests that these quiescent cell subpopulations in Multiple Myeloma are resistant to many therapies and so targeting these cells should be a priority for therapeutics.

In summary, this study shows for the first time that GSK343 appears to synergistically induce TRAIL responses Multiple Myeloma cells and responses are enhanced in 3D cell culture which is more representative of the tumour microenvironment. GSK343 appeared to lose its sensitising effect in TRAIL resistant RPMI 8226 cells with GSK343 +/−TRAIL having little effect on apoptosis induction, however NCI-H 929 retained sensitivity to dual treatment. GSK343 also had little effect on quiescent cells, analogous with the cancer stem cell phenotype which are drug resistant dormant cells responsible for tumour repopulation and Multiple Myeloma relapse.

## Notes

### Competing Interest Statement

The authors have declared no competing interest.

## References

Adomako A, Calvo V, Biran N, Osman K, Chari A, Paton JC, Paton AW, Moore K, Schewe DM, Aguirre-Ghiso JA. (2015) Identification of markers that functionally define a quiescent multiple myeloma cell sub-population surviving bortezomib treatment. BMC Cancer. 30, 444. doi: 10.1186/s12885-015-1460-1.

Alzrigat M, Párraga AA, Agarwal P, Zureigat H, Österborg A, Nahi H, Ma A, Jin J, Nilsson K, Öberg F, Kalushkova A, Jernberg-Wiklund H. (2017) EZH2 inhibition in multiple myeloma downregulates myeloma associated oncogenes and upregulates microRNAs with potential tumor suppressor functions. Oncotarget. 8, 10213–10224. doi: 10.18632/oncotarget.14378.

Agarwal P, Alzrigat M, Párraga AA, Enroth S, Singh U, Ungerstedt J, Österborg A, Brown PJ, Ma A, Jin J, Nilsson K, Öberg F, Kalushkova A, Jernberg-Wiklund H. (2016) Genome-wide profiling of histone H3 lysine 27 and lysine 4 trimethylation in multiple myeloma reveals the importance of Polycomb gene targeting and highlights EZH2 as a potential therapeutic target. Oncotarget. 7, 6809–23. doi: 10.18632/oncotarget.6843.

Arhoma A, Chantry AD, Haywood-Small SL, Cross NA. (2017) SAHA-induced TRAIL-sensitisation of Multiple Myeloma cells is enhanced in 3D cell culture. Exp Cell Res. 360, 226–235. doi: 10.1016/j.yexcr.2017.09.012.

Asangani IA, Ateeq B, Cao Q, Dodson L, Pandhi M, Kunju LP, Mehra R, Lonigro RJ, Siddiqui J, Palanisamy N, Wu YM, Cao X, Kim JH, Zhao M, Qin ZS, Iyer MK, Maher CA, Kumar-Sinha C, Varambally S, Chinnaiyan AM. (2013) Characterization of the EZH2-MMSET histone methyltransferase regulatory axis in cancer. Mol Cell. 49, 80–93. doi: 10.1016/j.molcel.2012.10.008.

Batie M, Frost J, Frost M, Wilson JW, Schofield P, Rocha S. (2019) Hypoxia induces rapid changes to histone methylation and reprograms chromatin. Science. 363, 1222–1226. doi: 10.1126/science.aau5870.

Benoit YD, Laursen KB, Witherspoon MS, Lipkin SM, Gudas LJ. (2013) Inhibition of PRC2 histone methyltransferase activity increases TRAIL-mediated apoptosis sensitivity in human colon cancer cells. J Cell Physiol. 228, 764–72. doi: 10.1002/jcp.24224.

Braun FK, Mathur R, Sehgal L, Wilkie-Grantham R, Chandra J, Berkova Z, Samaniego F. (2015) Inhibition of methyltransferases accelerates degradation of cFLIP and sensitizes B-cell lymphoma cells to TRAIL-induced apoptosis. PLoS One. 10, e0117994. doi: 10.1371/journal.pone.0117994.

Chen X, Thakkar H, Tyan F, Gim S, Robinson H, Lee C, Pandey SK, Nwokorie C, Onwudiwe N, Srivastava RK. (2001) Constitutively active Akt is an important regulator of TRAIL sensitivity in prostate cancer. Oncogene. 20,6 073–83. doi: 10.1038/sj.onc.1204736.

Chen Z, Orlowski RZ, Wang M, Kwak L, McCarty N. (2014) Osteoblastic niche supports the growth of quiescent multiple myeloma cells. Blood. 23, 2204–8. doi: 10.1182/blood-2013-07-517136.

Croonquist PA, Van Ness B. (2005) The polycomb group protein enhancer of zeste homolog 2 (EZH 2) is an oncogene that influences myeloma cell growth and the mutant ras phenotype. Oncogene 24, 6269–80. doi: 10.1038/sj.onc.1208771.

Dimopoulos K, Søgaard Helbo A, Fibiger Munch-Petersen H, Sjö L, Christensen J, Sommer Kristensen L, Asmar F, Hermansen NEU, O’Connel C, Gimsing P, Liang G, Grønbaek K. (2018) Dual inhibition of DNMTs and EZH2 can overcome both intrinsic and acquired resistance of myeloma cells to IMiDs in a cereblon-independent manner. Mol Oncol. 12, 180–195. doi: 10.1002/1878-0261.12157.

Drelon C, Berthon A, Mathieu M, Ragazzon B, Kuick R, Tabbal H, Septier A, Rodriguez S, Batisse-Lignier M, Sahut-Barnola I, Dumontet T, Pointud JC, Lefrançois-Martinez AM, Baron S, Giordano TJ, Bertherat J, Martinez A, Val P. (2016) EZH2 is overexpressed in adrenocortical carcinoma and is associated with disease progression. Hum Mol Genet 25, 2789–2800. doi: 10.1093/hmg/ddw136.

Geng C, Hou J, Zhao Y, Ke X, Wang Z, Qiu L, Xi H, Wang F, Wei N, Liu Y, Yang S, Wei P, Zheng X, Huang Z, Zhu B, Chen WM. (2014) A multicenter, open-label phase II study of recombinant CPT (Circularly Permuted TRAIL) plus thalidomide in patients with relapsed and refractory multiple myeloma. Am J Hematol. 89, 1037–42. doi: 10.1002/ajh.23822.

Gobbi G, Masselli E, Micheloni C, Nouvenne A, Russo D, Santi P, Matteucci A, Cocco L, Vitale M, Mirandola P. (2010) Hypoxia-induced down-modulation of PKCepsilon promotes trail-mediated apoptosis of tumor cells. Int J Oncol. 37, 719–29. doi: 10.3892/ijo_00000721.

Herbst RS, Eckhardt SG, Kurzrock R, Ebbinghaus S, O’Dwyer PJ, Gordon MS, Novotny W, Goldwasser MA, Tohnya TM, Lum BL, Ashkenazi A, Jubb AM, Mendelson DS. (2010) Phase I dose-escalation study of recombinant human Apo2L/TRAIL, a dual proapoptotic receptor agonist, in patients with advanced cancer. J Clin Oncol. 2010 28, 2839–46. doi: 10.1200/JCO.2009.25.1991.

Kalushkova A, Fryknäs M, Lemaire M, Fristedt C, Agarwal P, Eriksson M, Deleu S, Atadja P, Osterborg A, Nilsson K, Vanderkerken K, Oberg F, Jernberg-Wiklund H. (2010) Polycomb target genes are silenced in multiple myeloma. PLoS One. 5, e11483. doi: 10.1371/journal.pone.0011483.

Krueger A, Baumann S, Krammer PH, Kirchhoff S. (2001) FLICE-inhibitory proteins: regulators of death receptor-mediated apoptosis. Mol Cell Biol. 21, 8247–54. doi: 10.1128/MCB.21.24.8247-8254.2001.

Lawson MA, McDonald MM, Kovacic N, Hua Khoo W, Terry RL, Down J, Kaplan W, Paton-Hough J, Fellows C, Pettitt JA, Neil Dear T, Van Valckenborgh E, Baldock PA, Rogers MJ, Eaton CL, Vanderkerken K, Pettit AR, Quinn JM, Zannettino AC, Phan TG, Croucher PI. (2015) Osteoclasts control reactivation of dormant myeloma cells by remodelling the endosteal niche. Nat Commun. 3, 8983. doi: 10.1038/ncomms9983.

Leng Y, Hou J, Jin J, Zhang M, Ke X, Jiang B, Pan L, Yang L, Zhou F, Wang J, Wang Z, Liu L, Li W, Shen Z, Qiu L, Chang N, Li J, Liu J, Pang H, Meng H, Wei P, Jiang H, Liu Y, Zheng X, Yang S, Chen W. (2017) Circularly permuted TRAIL plus thalidomide and dexamethasone versus thalidomide and dexamethasone for relapsed/refractory multiple myeloma: a phase 2 study. Cancer Chemother Pharmacol. 79, 1141–1149. doi: 10.1007/s00280-017-3310-0.

Lo Sardo F, Pulito C, Sacconi A, Korita E, Sudol M, Strano S, Blandino G. (2021) YAP/TAZ and EZH2 synergize to impair tumor suppressor activity of TGFBR2 in non-small cell lung cancer. Cancer Lett. 500, 51–63. doi: 10.1016/j.canlet.2020.11.037.

Morgan GJ, Walker BA, Davies FE. (2012) The genetic architecture of multiple myeloma. Nat Rev Cancer. 12, 335–48. doi: 10.1038/nrc3257.

Nylund P, Atienza Párraga A, Haglöf J, De Bruyne E, Menu E, Garrido-Zabala B, Ma A, Jin J, Öberg F, Vanderkerken K, Kalushkova A, Jernberg-Wiklund H. (2021) A distinct metabolic response characterizes sensitivity to EZH2 inhibition in multiple myeloma. Cell Death Dis. 12, 167. doi: 10.1038/s41419-021-03447-8.

Pan G, O’Rourke K, Chinnaiyan AM, Gentz R, Ebner R, Ni J, Dixit VM. (1997) The receptor for the cytotoxic ligand TRAIL. Science. 276, 111–3. doi: 10.1126/science.276.5309.111.

Phillips KL, Wright N, McDermott E, Cross NA. (2019) TRAIL responses are enhanced by nuclear export inhibition in osteosarcoma. Biochem Biophys Res Commun. 517, 383–389. doi: 10.1016/j.bbrc.2019.07.047.

Popovic R, Martinez-Garcia E, Giannopoulou EG, Zhang Q, Zhang Q, Ezponda T, Shah MY, Zheng Y, Will CM, Small EC, Hua Y, Bulic M, Jiang Y, Carrara M, Calogero RA, Kath WL, Kelleher NL, Wang JP, Elemento O, Licht JD. (2014) Histone methyltransferase MMSET/NSD2 alters EZH2 binding and reprograms the myeloma epigenome through global and focal changes in H3K36 and H3K27 methylation. PLoS Genet. 10, e1004566. doi: 10.1371/journal.pgen.1004566.

Rajkumar SV, Kumar S. (2020) Multiple myeloma current treatment algorithms. Blood Cancer J. 10, 94. doi: 10.1038/s41408-020-00359-2.

Rajkumar SV. (2020) Multiple myeloma: 2020 update on diagnosis, risk-stratification and management. Am J Hematol. 95, 548–567. doi: 10.1002/ajh.25791.

Sayers, T and Cross, NA (2014) Triggering death receptors as a means of inducing tumoricidal activity. Chapter 8 in: Tumour Immunology and Immunotherapy. Oxford University Press. ISBN-13: 9780199676866

Schewe DM, Aguirre-Ghiso JA. (2009) Inhibition of eIF2alpha dephosphorylation maximizes bortezomib efficiency and eliminates quiescent multiple myeloma cells surviving proteasome inhibitor therapy. Cancer Res. 69, 1545–52. doi: 10.1158/0008-5472.CAN-08-3858.

Singh AK, Verma A, Singh A, Arya RK, Maheshwari S, Chaturvedi P, Nengroo MA, Saini KK, Vishwakarma AL, Singh K, Sarkar J, Datta D. (2021) Salinomycin inhibits epigenetic modulator EZH2 to enhance death receptors in colon cancer stem cells. Epigenetics. 16, 144–161. doi: 10.1080/15592294.2020.1789270.

Varambally S, Dhanasekaran SM, Zhou M, Barrette TR, Kumar-Sinha C, Sanda MG, Ghosh D, Pienta KJ, Sewalt RG, Otte AP, Rubin MA, Chinnaiyan AM. (2002) The polycomb group protein EZH2 is involved in progression of prostate cancer. Nature. 419, 624–9. doi: 10.1038/nature01075.

Vitovski S, Chantry AD, Lawson MA, Croucher PI. (2012) Targeting tumour-initiating cells with TRAIL based combination therapy ensures complete and lasting eradication of multiple myeloma tumours in vivo. PLoS One. 7,e35830. doi: 10.1371/journal.pone.0035830.

Wouters BG, Koritzinsky M. (2008) Hypoxia signalling through mTOR and the unfolded protein response in cancer. Nat Rev Cancer. 8, 851–64. doi: 10.1038/nrc2501.

Verma SK, Tian X, LaFrance LV, Duquenne C, Suarez DP, Newlander KA, Romeril SP, Burgess JL, Grant SW, Brackley JA, Graves AP, Scherzer DA, Shu A, Thompson C, Ott HM, Aller GS, Machutta CA, Diaz E, Jiang Y, Johnson NW, Knight SD, Kruger RG, McCabe MT, Dhanak D, Tummino PJ, Creasy CL, Miller WH. (2012) Identification of Potent, Selective, Cell-Active Inhibitors of the Histone Lysine Methyltransferase EZH2. ACS Med Chem Lett 3, 1091–6. doi: 10.1021/ml3003346.

Yap TA, Winter JN, Giulino-Roth L, Longley J, Lopez J, Michot JM, Leonard JP, Ribrag V, McCabe MT, Creasy CL, Stern M, Pene Dumitrescu T, Wang X, Frey S, Carver J, Horner T, Oh C, Khaled A, Dhar A, Johnson PWM. (2019) Phase I Study of the Novel Enhancer of Zeste Homolog 2 (EZH2) Inhibitor GSK2816126 in Patients with Advanced Hematologic and Solid Tumors. Clin Cancer Res. 25, 7331–7339. doi: 10.1158/1078-0432.CCR-18-4121.

Zhang Q, Padi SK, Tindall DJ, Guo B. (2014) Polycomb protein EZH2 suppresses apoptosis by silencing the proapoptotic miR-31. Cell Death Dis. 5, e1486. doi: 10.1038/cddis.2014.454.

